# Local FK506 delivery induces osteogenesis in *in vivo* rat bone defect and rabbit spine fusion models

**DOI:** 10.1101/2024.03.08.584163

**Authors:** Julia Andraca Harrer, Travis M. Fulton, Sreedhara Sangadala, Jarred Kaiser, Emily J. Devereaux, Colleen Oliver, Steven M. Presciutti, Scott D. Boden, Nick J. Willett

**Author notes:** Correspondence (NJW). These authors have contributed equally to this work and share first authorship (Julia Andraca Harrer, Travis M. Fulton).

## Abstract

Bone grafting procedures are commonly used for the repair, regeneration, and fusion of bones in in a wide range of orthopaedic surgeries, including large bone defects and spine fusion procedures. Autografts are the clinical gold standard, though recombinant human bone morphogenetic proteins (rhBMPs) are often used, particularly in difficult clinical situations. However, treatment with rhBMPs can have off-target effects and significantly increase surgical costs, adding to patients’ already high economic and mental burden. Recent studies have identified that FDA-approved immunosuppressant drug, FK506 (Tacrolimus), can also activate the BMP pathway by binding to its inhibitors. This study tested the hypothesis that FK506, as a standalone treatment, could induce osteogenic differentiation of human mesenchymal stromal cells (hMSCs), as well as functional bone formation in a rat segmental bone defect model and rabbit spinal fusion model. FK506 potentiated the effect of low dose BMP-2 to enhance osteogenic differentiation and mineralization of hMSCs *in vitro*. Standalone treatment with FK506 delivered on a collagen sponge, produced consistent bone bridging of a rat critically-sized femoral defect with functional mechanical properties comparable to naïve bone. In a rabbit single level posterolateral spine fusion model, treatment with FK506 delivered on a collagen sponge successfully fused the L5-L6 vertebrae at rates comparable to rhBMP-2 treatment. These data demonstrate the ability of FK506 to induce bone formation in human cells and two challenging *in vivo* models, and indicate FK506 can be utilized either as a standalone treatment or in conjunction with rhBMP to treat a variety of spine disorders.

**Graphical Abstract:** 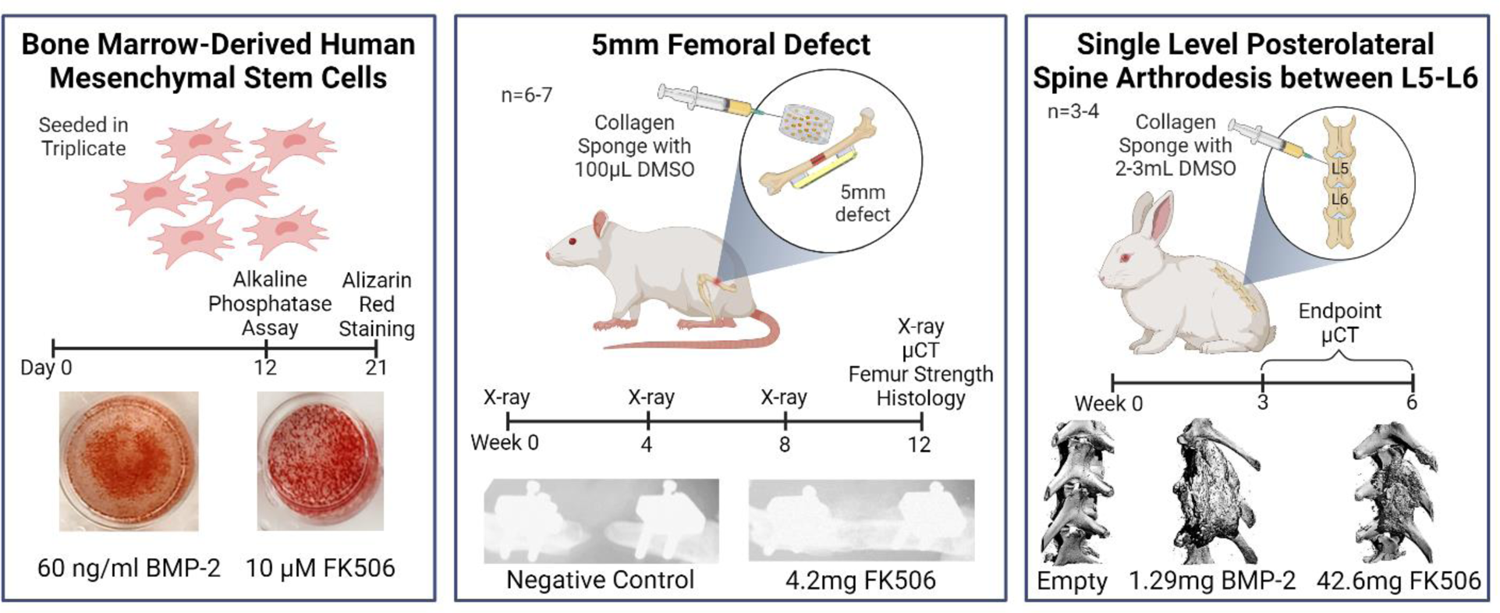

## Introduction

Bone grafting procedures are commonly used in a wide variety of orthopaedic procedures ranging from regeneration of bone defect procedures to fusion of joints, such as during spinal fusion procedures. Spinal fusion is used to treat a variety of spine disorders including tumors, deformities, and traumatic injuries. Success rates reported for these procedures vary greatly, with non-union rates ranging from 1.1%-43.3%, and can take up to 3 years to reach union ^1–4^. The incidence of spinal fusion and refusion procedures has increased drastically in recent years, with a 137% and 187% increase from 1998 to 2013, respectively ^5,6^. Historically, the most common method of achieving spinal fusion is through an autologous bone graft treatment, which has potential limitations and complications including increased infection rates, donor site pain and morbidity, limited tissue availability, and revision surgeries ^7–11^. In 2002, recombinant human Bone Morphogenetic Protein-2 (rhBMP-2) was FDA approved for use in spine surgeries, and has been increasingly employed in spine fusions as an alternative to autograft procedures ^12,13^. However, surgeries using rhBMP-2 have increased rates of inflammation, radiculitis, bone resorption and ectopic bone formation, as well as the possibility of developing BMP-2 neutralizing antibodies ^12,14,15^. Additionally, initial rhBMP-2 surgeries increase surgical costs by $15,000 on average ^13,16^. A combination of cost, complications, and transportation/storage difficulties has led to an increased interest in the use of alternate approaches, such as osteoinductive small molecule treatment, to replace rhBMP-2’s role in bone regeneration ^17^.

FK506, or Tacrolimus, is an FDA approved small molecule that is currently used as an immunosuppressant for organ transplant patients ^18,19^. FK506 has also been shown to enhance BMP-2 activity and osteogenic cell differentiation by binding to BMP inhibitor FKBP12 and inhibiting calcineurin activity ^20,21^. *In vitro* studies have demonstrated that FK506 promotes osteogenic differentiation in multiple rodent cell lines, including murine C2C12 myoblasts, murine MC3T3 pre-osteoblasts, and rat bone marrow mesenchymal stromal cells ^22,23^. Most *in vivo* studies investigating the potential of FK506 in bone regeneration have used the drug as an immunosuppressant to supplement other treatments. In a rat ectopic bone formation model, local injection of FK506 along with BMP-2 or a BMP-2-expressing recombinant adenoviral vector improved bone formation ^24,25^. In other ectopic bone formation models, the supplementation of isograft, allograft, or demineralized bone matrix implantations with FK5606 demonstrated that FK506 injection significantly enhanced bone formation ^26,27^. In orthotopic bone defect models, systemic FK506, as an immunosuppressant, demonstrated enhanced the osteogenic effects of local bone regeneration, including demineralized bone matrix and BMP-2 expressing adenovirus ^28,29^. While these data have demonstrated clear potential of FK506 as a supplemental treatment during bone regeneration procedures, the utility and potential of local, standalone treatment of FK506 for clinically relevant procedures, such as bone defect repair and spinal fusion surgeries, has yet to be investigated.

In previous studies, we evaluated the ability of FK506 as a standalone treatment to induce osteogenic differentiation in murine C2C12 cells and an ectopic bone formation model in rats ^30^. Culturing cells with FK506 induced alkaline phosphatase (ALP) activity at a comparable level to BMP-2. Additionally, cells treated with FK506 showed increased pSMAD, ID1, and TIEG1 levels, indicating activation of BMP and TGF-β pathways. FK506-loaded collagen sponges implanted subcutaneously in rats induced bone volume and mineralization levels comparable to bone volume and mineralization induced by BMP-2. These data demonstrate the potential of FK506 as a standalone treatment and alternative to BMP-2 for bone healing and spinal fusion procedures.

The objective of this current study was to evaluate the effects of standalone FK506 treatment on osteogenic differentiation in human mesenchymal stromal cells and as a locally delivered treatment for both segmental bone defects and spinal fusion. We hypothesized that FK506 would induce bone formation in each model at a level comparable to that of BMP-2. This study assessed *in vitro* treatment of FK506 in human bone marrow-derived MSCs both in conjunction with BMP-2 and independently. We also assessed standalone treatment of FK506 in an established rat femoral defect model and dose response studies in a rabbit single level posterolateral spine fusion model to evaluate the efficacy of FK506 as a potential treatment for bone defect and spinal disorders.

## Results

### FK506 Induced Mineralization in Adult Human Mesenchymal Stromal Cells

Adult human mesenchymal stromal cells (hMSCs) were cultured with hMSC growth media, osteogenic media, or osteogenic media with various concentrations of BMP-2 or FK506. Induction of early osteogenic differentiation was measured through alkaline phosphatase (ALP) expression at day 12 (Fig. 1A). All doses of BMP-2 treatment resulted in significantly higher ALP activity compared to the growth media and osteogenic media (0 ng BMP-2) controls (Supplementary Table S1). Similarly, all doses of FK506 treatment resulted in significantly higher ALP activity compared to the growth media and osteogenic media (0 µM FK506) controls. Both treatment with 160 ng BMP-2 and 240 ng BMP-2 resulted in significantly higher ALP activity compared to 10 µM FK506 (160 ng BMP-2: 5.857 nmol/µg ± 0.02443 nmol/µg and 240 ng BMP-2: 8.087 nmol/µg ± 0.04737 nmol/µg vs. 10 µM FK506: 4.250 nmol/µg ± 0.02259 nmol/µg, p < 0.001).

**Figure 1.**
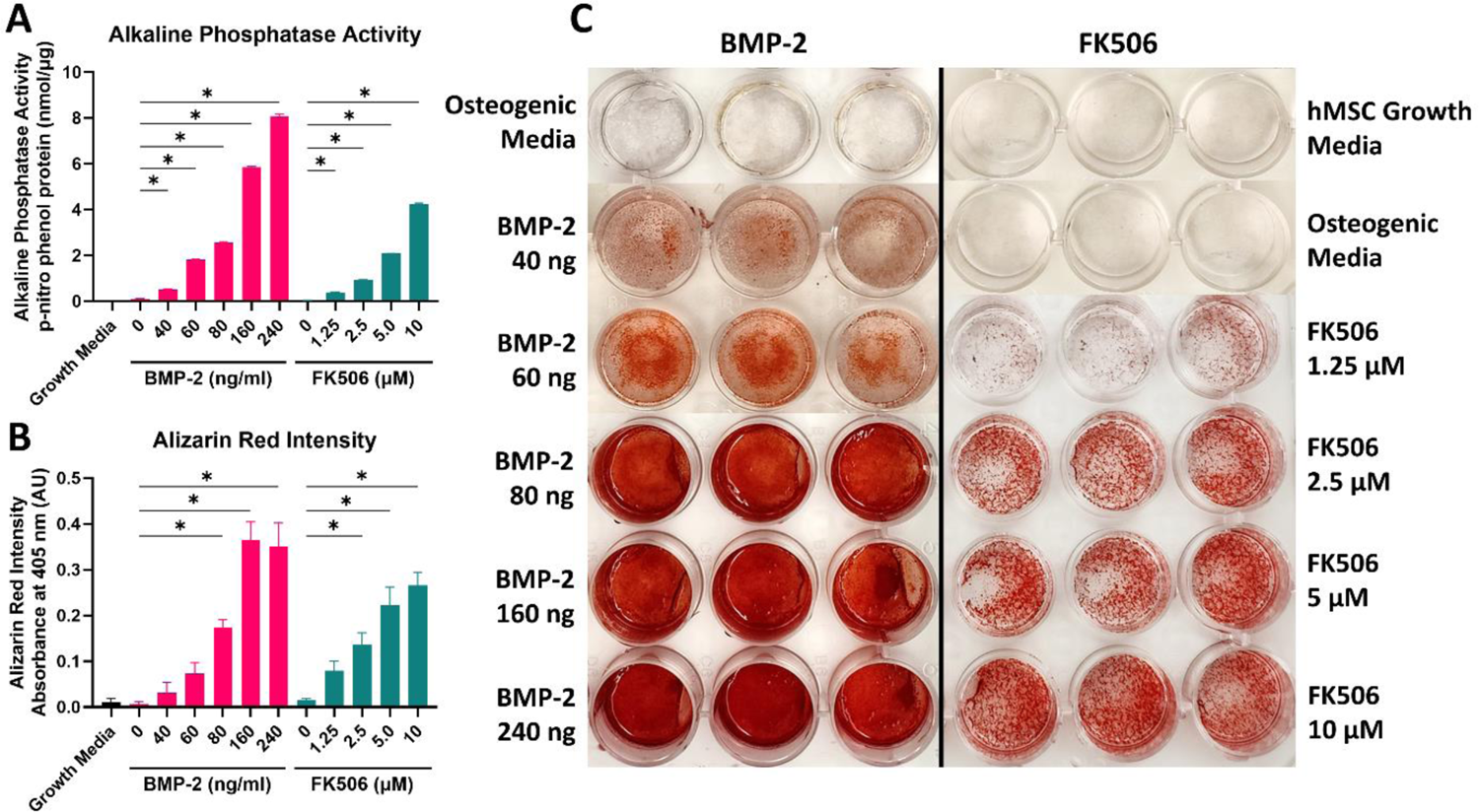
Human mesenchymal stromal cells showed increased osteogenic differentiation in response to FK506. Quantification of alkaline phosphatase activity (A) and Alizarin red stain for calcium mineralization (B). Alizarin red staining (C). One-way ANOVA with Tukey’s multiple comparisons.

Induction of calcium mineralization of hMSCs at day 21 in response to increasing doses of BMP-2 and FK506 was analyzed by staining with alizarin red (Fig. 1C). Treatment with 80 ng, 160 ng, and 240 ng BMP-2 and 2.5 µM, 5.0 µM, and 10 µM FK506 were significantly increased compared to growth media and osteogenic media controls (Supplementary Table S2). Treatment with 160 ng and 240 ng BMP-2 was significantly higher than all doses of FK506 (160 ng BMP-2: 0.3643 AU ± 0.02325 AU vs 10 µM FK506: 0.2660 AU ± 0.01652 AU, p = 0.0094, and 240 ng BMP-2: 0.3507 AU ± 0.02973 AU vs 10 µM FK506: 0.2660 AU ± 0.01652 AU, p = 0.0372).

Treatment with 80 ng BMP-2 was not significantly different from treatment with and 2.5 µM and 5.0 µM FK506, but was significantly lower than treatment with 10 µM FK506 (80 ng BMP-2: 0.1733 AU ± 0.01033 AU vs. 2.5 µM FK506: 0.1373 AU ± 0.01443 AU, p = 0.8963, 5.0 µM FK506: 0.2223 AU ± 0.02310 AU, p = 0.5864, and 10 µM FK506: 0.2660 AU ± 0.01652 AU, p = 0.0168).

### Local Delivery of FK506 Bridges Critically-Sized Defects in Rat Femurs

A critically-sized 5 mm bone defect was created in 14 rat femurs to compare the treatment effects of 4.2 mg of FK506 in DMSO to a negative control DMSO delivery on a collagen sponge. Radiographs were obtained immediately after surgery and at four-week intervals until the study endpoint at Week 12 (Fig. 2). Radiographs were evaluated by three blinded scorers; 7/7 femurs bridged in the FK506 treatment group and 0/6 in the negative control group. In the FK506 group, radiographic evidence of bridging was noted as early as 4 weeks post-surgery.

**Figure 2.**
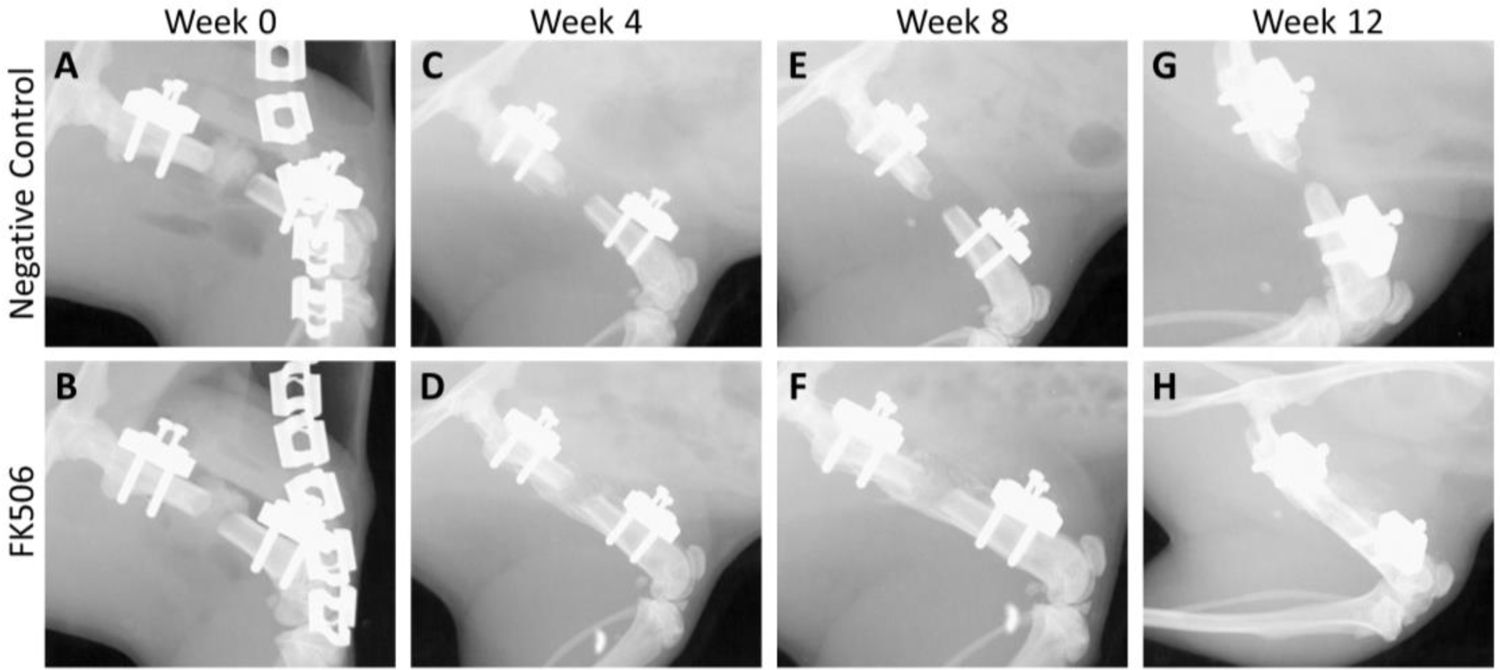
Representative radiographs of defect site at (A,B) post-operation, (C,D) 4 weeks, (E,F) 8 weeks, and (G,H) 12 weeks.

All femurs were resected for micro-computed tomography (µCT) analysis and mechanical testing. The defect region showed substantially greater bone volume in the FK506 treatment group compared to the negative control group (56.63 mm^3^ ± 5.007 vs. 5.4 mm^3^ ± 0.7191, p < 0.001) and bone mineral density was lower in FK506 treated defects compared to those from negative control femurs (444.48 mgHA/cm^3^ ± 42.74 vs. 651.96 mgHA/cm^3^ ± 7.19, p < 0.001) (Fig. 3 A,B, E, F).

**Figure 3.**
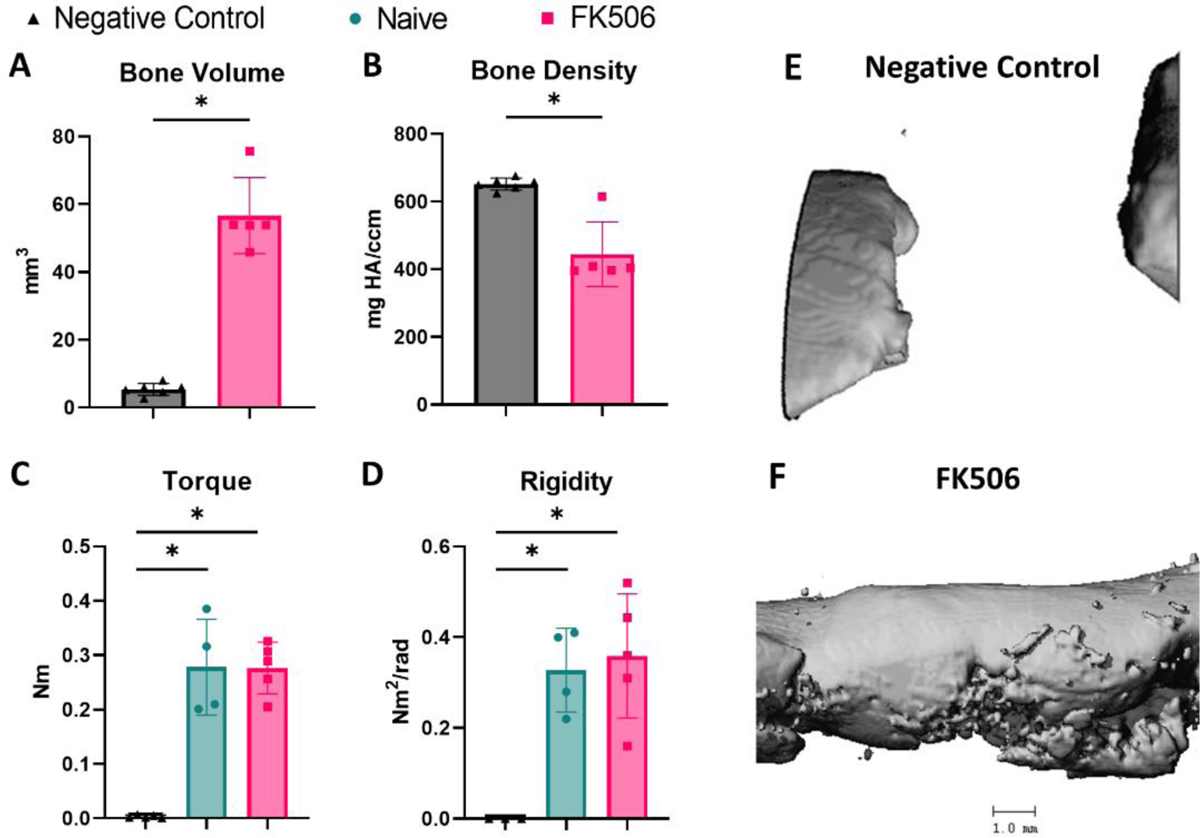
µCT analysis of bone volume and density showed increased bone volume in FK506 group compared to the negative control (A,B). Results of maximum torque to failure and rigidity mechanical testing demonstrated that FK506 treated femurs and naïve control femurs had no differences in mechanical properties, both were significantly higher than the negative control group (C,D). µCT 3D reconstruction of defect site at 12 weeks (E,F). * denotes p < 0.05, unpaired t test (A,B) and one-way ANOVA with Tukey’s multiple comparisons (C,D).

When evaluating mechanical properties of the femurs, contralateral limbs were used as naïve controls. Naïve, FK506 treated, and negative control femora were evaluated for maximum torque to failure and rigidity. FK506 treated femurs were found to have no significant difference in maximum torque to failure compared to naïve femurs (0.2764 Nmm ± 0.02118 vs. 0.2780 Nmm ± 0.0442, p = 0.9989) and FK506 treated femurs also had no significant differences in rigidity compared to the naïve femurs (0.3586 Nmm^2^/rad ± 0.06124 vs. 0.3275 Nmm^2^/rad ± 0.04644, p = 0.9006) (Fig. 3 C,D). The negative control group was significantly different from both the FK506 treated femurs and the naïve control groups for both measurements (Torque: 0.0034 ± 0.001122 Nmm, p < 0.001, Rigidity: 0 Nmm^2^/rad, negative control vs. FK506 p = 0.0031, negative control vs. naïve control p = 0.0073).

Following µCT analysis, rat femora were decalcified, fixed, sectioned, and stained with Hematoxylin and Eosin (H&E), Goldner’s Trichrome, and Safranin-O/Fast-green. Femora in the FK506 treatment group were prepped for histology immediately after µCT, while femora in the negative control group were prepped for histology following mechanical testing. In the negative control group, no bridging was visible, with some fibrous tissue forming between bone caps on either end of the defect (Fig. 4 A-F). Robust bridging was visible in FK506-treated femora, with cortical bone fully surrounding the trabecular bone forming within (Fig. 4 G-L). No cartilage was seen bridging or around the defect region in either group.

**Figure 4.**
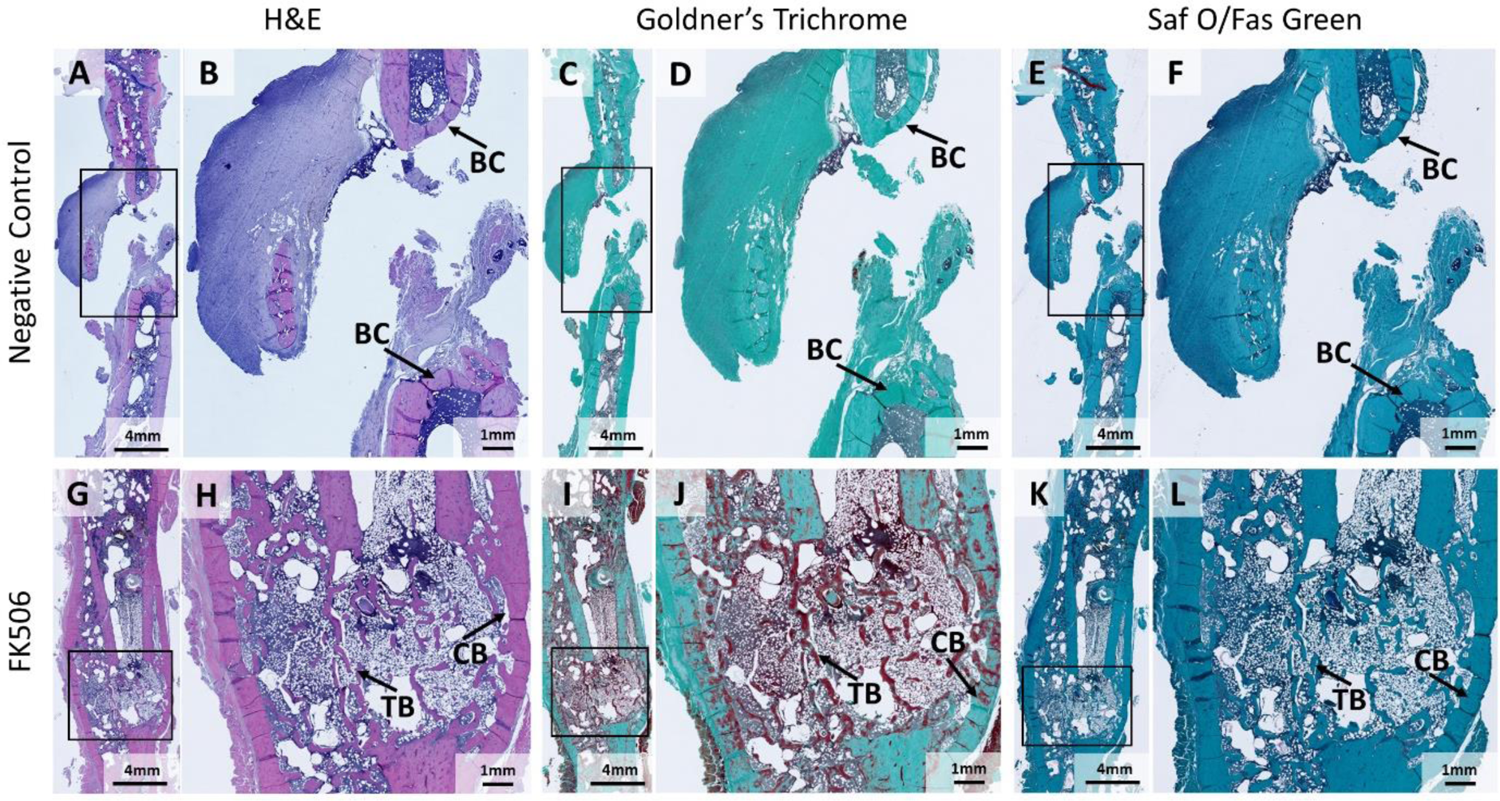
Hematoxylin and Eosin (A,B, G,H), Goldner’s Trichrome (C,D,I,J), Safranin-O (E,F,K,L) staining of negative control defect site and FK506 defect site respectively at 12 weeks. Stained longitudinal sections of defect site, box denotes defect site (A,C,E,G,I,K). Enlarged view of defect sites, arrows identify trabecular bone growth (TB), cortical bone growth (CB), and bone cap formation (BC) (B,D,F,H,J,L). Both trabecular and cortical bone bridge FK506 femora, while bone caps indicate non-union in negative control group.

### Local Delivery of FK506 Induces Spinal Fusion between Rabbit Lumbar Vertebrae

Ten New Zealand White rabbits underwent a single level posterolateral spine arthrodesis procedure between L5-L6 to compare BMP-2 treatment to varying doses of FK506. All treatments were delivered by first injecting the treatment solution onto the collagen sponge and then implanting the sponge between the L5-L6 vertebrae. Bilateral procedures were performed for two treatment groups, with 1.29 mg BMP-2 on one side and 145 mg FK506 on the other side (n=4). Unilateral procedures were performed for the 64.3 mg FK506 and 42.6 mg FK506 treatment groups (n=3), with no treatment on the other side. Rabbits with higher FK506 dosage levels (4/4 rabbits with 145 mg FK506, 1/3 with 64.3 mg FK506, 0/3 with 42.3 mg FK506) lost over 25% of their body weight over the course of the study compared to their baseline weight recorded at time of surgery; these animals were euthanized at the timepoint of these measurements which ranged from 3-6 weeks post-surgery. Following euthanasia, animals were examined, and it was noted that all rabbits with weight loss appeared to have large volumes of hard, impacted food in the stomach and distended intestines.

Rabbits from 1.29 mg BMP, 145 mg FK506, 64.3 mg FK506, and 42.6 mg FK506 groups showed varied amounts of bone formation and spine fusion. Rabbits in the 145 mg FK506 treatment group had bone formation in 3/4 samples and rabbits in the 1.29 mg BMP-2, 64.3 mg FK506, and 42.6 mg FK506 groups had bone formation in all animals (Table 1). Successful fusion between L5-L6 vertebrae was evaluated by 2 blinded orthopaedic surgeon scorers. The 1.29 mg BMP group had 3/4 successful fusions, 145 mg FK506 had 1/4, 64.3 mg FK506 had 1/3, and 42.6 mg FK506 had 2/3 successful fusions (Table 1). µCT scans were used to evaluate bone volume and mineralization (Fig. 5). All treatment groups except the 64.3 mg FK506 group had significantly greater bone volume compared to the no treatment group (No treatment: 0.06437 mm^3^ ± 0.02296 vs. BMP-2: 1506.01 mm^3^ ± 210.8, p < 0.001, 145 mg FK506: 671.09 mm^3^ ± 264.0, p = 0.0238, 64.3 mg FK506: 582.53 mm^3^ ± 132.4, p = 0.0616, 42.6 mg FK506: 684.4 mm^3^ ± 79.78, p = 0.0207). Bone volume due to BMP treatment was significantly greater than all treatment groups, however the BMP-induced bone formation resulted in notable ectopic bone formation often bridging well past a single level fusion (BMP-2: 1506.01 mm^3^ ± 210.8 vs. 145 mg FK506: 671.09 mm^3^ ± 264.0, p = 0.0137, 64.3 mg FK506: 582.53 mm^3^ ± 132.4, p = 0.0062, 42.6 mg FK506: 684.4 mm^3^ ± 79.78, p = 0.0155, no treatment: 0.06437 mm^3^ ± 0.02296, p < 0.001). There was no significant difference in bone mineral density between groups (BMP: 446.0 mgHA/cm^3^ ± 22.30, 145 mg FK506: 398.9 mgHA/cm^3^ ± 8.565, 64.3 mg FK506: 426.3 mgHA/cm^3^ ± 8.004, 42.6 mg FK506: 430.0 mgHA/cm^3^ ± 15.19).

**Figure 5:**
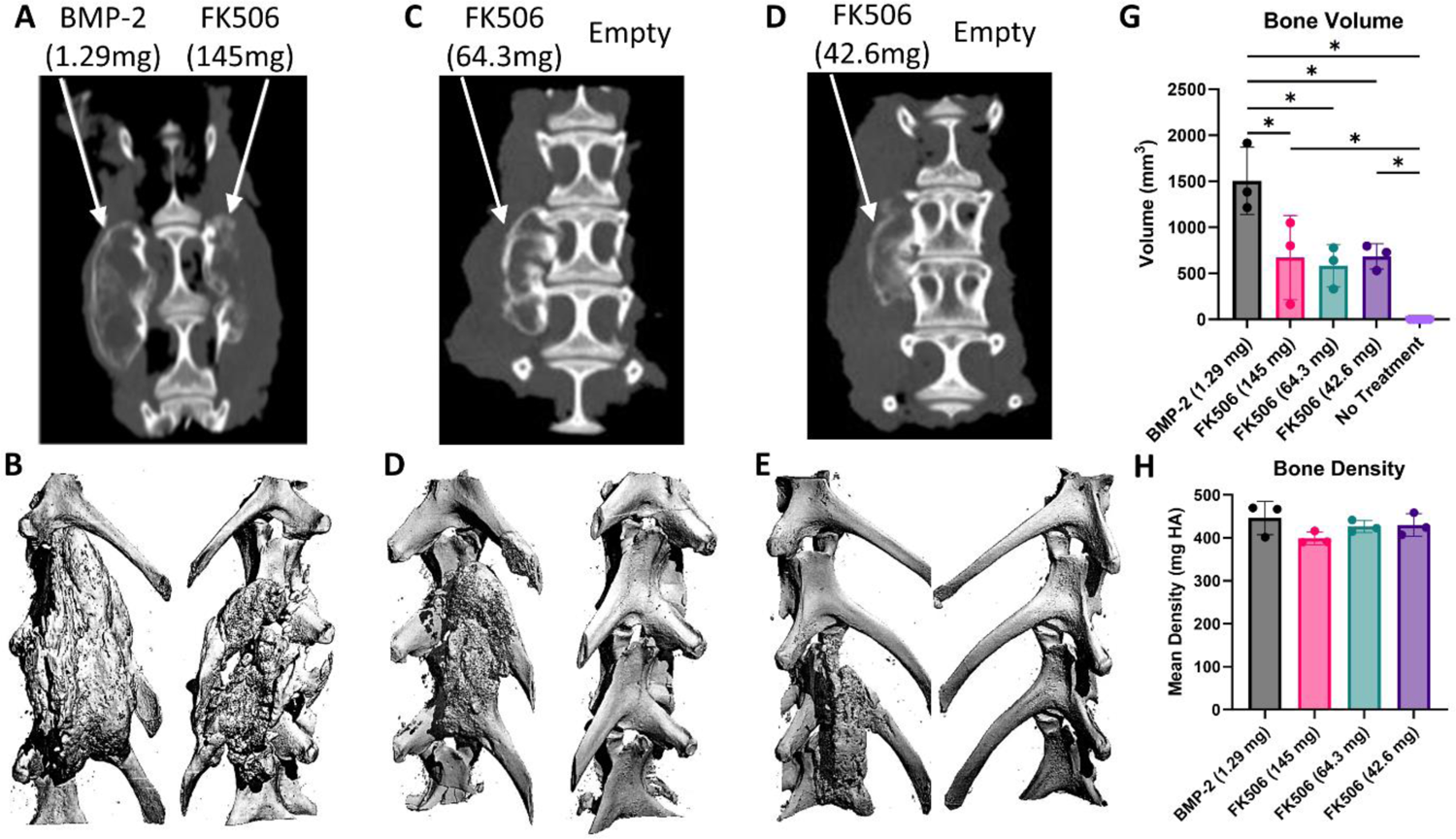
µCT analysis of bone volume and density. µCT slices (A,C,E) and 3-dimensional reconstruction (B,D,F) showing bone formation. µCT analysis of bone volume showed significant differences between the no treatment group and all treatment groups except 64.3 mg FK506 treatment group (G). There was no significant difference in bone mineral density in all groups (H). * denotes p < 0.05, one-way ANOVA with Tukey’s multiple comparisons.

**Table 1:** Evaluation of bone formation and fusion. Blinded scorers evaluated whether bone formation and fusion were noted in each rabbit. Both scorers were orthopaedic spinal surgeons.

| Treatment | Bone Formation | Fused |
| --- | --- | --- |
| rh-BMP-2 (1.29mg) | 4/4 | 3/4 |
| FK506 (145mg) | 3/4 | 1/4 |
| FK506 (64.3mg) | 3/3 | 1/3 |
| FK506 (42.6mg) | 3/3 | 2/3 |

## Discussion

The incidence of bone grafting procedures has consistently increased in recent years; however, complication rates have not improved, underscoring the need to improve the current method of treatment ^5,6,33^. rhBMP-2 is FDA approved for spine fusion and has been increasingly used as an alternative to autografts, however both autografts and BMPs have associated complications. rhBMP-2 has reported adverse side effects that can occur including infection, inflammation, ectopic bone formation, bone resorption, neurological effects, radiculitis, and negative impacts to several other organ systems in the body ^15,34–37^. In addition to these effects, BMP-induced bone formation at supra-physiological doses can produce thin and wispy bone that has cyst-like properties filled with adipose tissue and low mechanical strength ^38,39^. As an alternative to using supraphysiologic doses of BMP, we investigated the potential to repurpose FK506, an FDA-approved immunosuppressant drug, to induce osteogenic differentiation and bone formation in multiple rigorous pre-clinical bone regeneration and fusion models. Our previous research demonstrated the ability of FK506 to induce bone formation as a standalone treatment by targeting the BMP pathway ^30^. Here we hypothesized that FK506 would induce mineralization in human mesenchymal stromal cells and bone formation in two rigorous *in vivo* models at comparable levels to rhBMP-2.

The clinical translation pathway of an osteoinductive therapy or bone graft alternative has a well-established series of pre-clinical milestones and models used to show therapeutic efficacy and safety. These include demonstrating efficacy in human cells in vitro, efficacy in small animal ectopic and orthotopic bone regeneration models, and then scaling to large animal orthotopic models (rabbits and then non-human primates). Here we assessed mineralization and osteogenic differentiation in hMSCs treated with increasing doses of FK506 and BMP-2 to assess responsiveness in human cells. Treatment with FK506 enhanced both mineralization and markers of osteoblastic differentiation in hMSCs, suggesting that FK506 could be a viable treatment either as a standalone treatment or to supplement lower BMP dosages in patients, potentially reducing adverse side effects associated with supraphysiologic doses of BMP. Osteogenic differentiation resulting from standalone FK506 treatment is consistent with other cell culture studies. In rat derived MSCs and human gingival derived stem cells that showed increased alizarin red staining, alkaline phosphatase activity, osteocalcin, and pSmad1/5 expression with standalone FK506 treatment ^22,40–42^. Gene expression analysis in a human osteoblast cell line treated with FK506 showed an increase in TGF-β1 receptor and Smad2 expression, along with EGF receptor, MMP2, biglycan, osteonectin, and collagen types III and XII ^43^. This indicates FK506 may play a role in osteogenesis through extracellular matrix formation and remodeling, osteoblast differentiation, and mineralization through TGF-β signaling ^44^. Examination of changes in miRNA expression following FK506-induced osteogenic differentiation demonstrated upregulation of Smad5, Jagged 1, and MAPK9 and downregulation of Smad7 and other negative regulators of osteogenic differentiation ^45^. These pathways can be downstream of both BMP signaling and calcineurin/FKBP12 signaling, as demonstrated in our previous studies ^30^. Future research could further discern the direct mechanism of action and to uncover any crosstalk between the different pathways activated by FK506.

Bone grafting procedures and osteoinductive therapies are commonly used in trauma injury procedures and segmental bone defect repair. We evaluated the ability of FK506 to induce bone formation in a well-established rat segmental bone defect model. Both bridging and bone formation were significantly enhanced by treatment with FK506 as seen by radiographs, μCT, and histological assessment. In addition, the regenerated bone from the FK506 treatment had similar properties to naïve bone, indicating functional restoration of the femur. While these data are generally consistent with previous literature, previous studies have shown mixed data in part due to a variety of local and systemic treatment modalities with FK506, as well as primarily using FK506 as an immunosuppressive supplement to other osteoinductive treatments rather than a standalone treatment. Previous studies using a rat bone defect model investigated the effects of systemic FK506 treatment in combination with a local treatment an adenovirus expressing BMP-2; this study showed that systemic FK506 treatment enhanced the bone regeneration from the local BMP-2 gene therapy ^28,29^. In another study in a critically-sized bone defect model, systemic delivery of FK506 was tested in combination with a local treatment of MSCs, BMP-2, or genetically modified MSCs expressing BMP-2. This study showed that FK506 delivery in the genetically modified MSCs, the FK506 treatment more than tripled the number of bridged bone defects, though FK506 did not impact healing outcomes in bone defects treated with rhBMP-2 or unmodified MSCs ^46^. While these studies all showed potential benefit of systemic FK506 delivery to potentiate a local osteoinductive treatment, these studies predominantly focused on FK506 as an indirect immunosuppressant, with doses at approximately 1 mg/kg rather than a direct osteoinductive treatment, indicating the dose may not be at a level needed to activate BMP signaling. As a standalone immunosuppressive treatment, systemic FK506 has been tested in a combined TA muscle trauma and endogenously healing tibial osteotomy model in a rat; this study showed that daily systemic delivery of FK506 alone at a 1 mg/kg dose improved mechanical properties of the tibia in the combined injury model, but did not affect mechanical properties in the isolated osteotomy model ^47^. This study showed that standalone systemic FK506 treatment could improve bone formation and healing, similar to the results seen in our study. The differences seen in the above studies could be due to the other treatments used in combination with FK506, the dose and delivery frequency of FK506, and the wide variety of carriers used for delivery. The various systemic delivery methods used may also have limited the effect on the bone healing pathway. In fact, there have been multiple studies, with contradictory results, investigating whether systemic delivery of FK506 for immunosuppression could induce osteopenia ^48–51^. Despite the variability in the method of delivery, multiple studies have demonstrated the potential for systemic FK506 treatment to induce local bone regeneration in a variety of bone defect models.

Local delivery of FK506 has been tested in various bone healing models. In rodent fracture and osteotomy models, local delivery through daily intramuscular injections or an osmotic pump at a 1 mg/kg or 10 mg/kg dose both showed no change in bone healing rate, with all fractures and defects bridging by weeks 3-4, regardless of FK506 dose ^52,53^. Both studies were performed in a sub-critical injury model which would already heal and did not have much range to see an effect of the FK506 treatment. One previous study investigated standalone local treatment of FK506 in a critically-sized, 7 mm rat calvarial defect model. In this study, 1 mg FK506 (approximately 3.7-4.0 mg/kg dose) loaded on a collagen hydrogel with a PLC/gelatin membrane was implanted into the calvarial defect and showed significantly increased bone formation compared to other groups^54^. This study is consistent with our findings that FK506 can be used as a standalone osteoinductive therapy in critically-sized bone defect and fusion procedures.

Another common bone grafting application, and the primary FDA approved clinical indication for rhBMP delivery, is in spine fusion procedures. The standard pre-clinical model is to use the rabbit spine fusion model before moving to a non-human primate model ^32^. We therefore aimed to test the efficacy of FK506 treatment in the rabbit spine fusion model. However, in previous studies investigating the effects of FK506 immunosuppression in rabbits, rabbits were shown to have a low toxicity threshold and elevated side effects in response to systemic FK506 levels including anorexia, weight loss, and renal complications ^55–57^. As such, multiple doses were tested to determine which dose increased bone formation while minimizing adverse effects. The dosing was scaled up from rat concentration to rabbit concentration based on metabolism scaling predictions as well as previous data from systemic rabbit toxicity data. Of the three doses tested, all rabbits in the highest dose category and one in the middle dose exhibited greater than 25% weight loss compared to baseline weight, impacted food in the stomach, and distended intestines. This indicates issues with gut secretions, mobility, or absorption ability; consistent with complications other groups have found specific to rabbit response to FK506 treatment ^55–57^. The rabbits treated with the lowest dose of FK506, however, did not show these side effects. Various studies in humans have detailed gastrointestinal effects of long-term delivery of FK506 as an immunosuppressant, including effects on absorption and intestinal barrier function ^19,58,59^. However, it is unclear how the effects of chronic, systemic, and repeated administration FK506 would compare to a single, localized high dose of FK506. This warrants further investigation into the effects of a high localized dose of FK506 as this treatment is scaled to larger pre-clinical and clinical models. Regardless of the side effects, local FK506 delivery produced bone formation at the spinal fusion site in all rabbits receiving the treatment. There was no difference in bone mineral density between rhBMP-2 and all FK506 doses, though rhBMP-2 treatment had statistically higher regenerated bone volume compared to all FK506 doses. However, only the lowest dose of FK506 resulted in comparable fusion rates to BMP-2. The fusion rates seen in the 42.6 mg FK506 and 1.29 mg BMP-2 groups (66% and 75%, respectively) are comparable to historical fusion rates achieved by the gold standard treatment, iliac crest autologous bone graft, in previous spinal fusion studies using this rabbit model. In the original study establishing the rabbit posterolateral spine arthrodesis model, autograft resulted in a 66% fusion rate at 6 weeks ^32^. Two reviews conducted comparing fusion rates in this model in 900 and 700 surgeries found that autografts resulted in 55% and 70% fusion rates, respectively ^60,61^. The fusion rates from the 42.6 mg FK506 treatment is within the range of the clinical gold standard and benchmark BMP-2 positive control. When looking at the 3D reconstructions of bone formation, treatment with BMP-2 results in bone formation beyond the single level fusion location up to the adjacent facet joint (Fig. 5 A,B). This highlights one of the more prominent issues with BMP treatment, ectopic bone formation, as fusion of an additional joint would limit the patient’s range of motion. Conversely, the bone formed by FK506 remained within the single joint space it was meant to fuse. This lack of ectopic bone formation resulting from FK506 treatment is corroborated by a study into a genetic disorder, Fibrodysplasia ossifans progressiva (FOP), which causes ectopic bone formation in soft tissues ^62^. When investigating the mechanism of action, it was found that FK506 did not enhance the responsiveness to endogenous BMPs, and therefore does not cause the ectopic bone formation seen in FOP ^62^. When viewed alongside the lack of ectopic bone formation in the rabbit spine fusion model (Fig. 5), this suggests that FK506 may reduce the potential complications related to ectopic bone formation observed with BMP-2.

While direct activation of FK506 in osteoprogenitor cells through activation of the BMP pathway may be a key mechanism of action for local FK506 induced bone formation, other actions of FK506 may also influence the bone regeneration and fusion. Complications due to inflammation is one of the major off-target effects of BMP-2, and there is potential that FK506 may suppress this response. FK506 is a potent immunosuppressant via suppression of T cell proliferation, which could be beneficial to the bone healing process. It is well known that chronic inflammation can result in poor bone healing outcomes ^63–65^. Specifically, imbalance in macrophage polarization, elevated myeloid-derived suppressor cells, and increased levels of pro-inflammatory cytokines have been implicated in reduced bone healing ^66–68^. The increased inflammatory response may inadvertently induce osteoclast activation which leads to increased bone resorption ^64^. FK506 functions through inhibiting calcineurin activity and thus preventing T cell proliferation and IL-2 expression ^20,21^. FK506 has also been shown to suppress osteoclast formation by decreasing NFATc1 activation, and has been successfully used in humans to induce bone formation ^69,70^. Further investigation into changes in cell populations and the cytokine profile after treatment with FK506 could further inform the role of FK506-induced immunosuppression in bone healing.

This study provides evidence that FK506 could be utilized as an alternative osteoinductive treatment for spine fusion and bone defect repair procedures; however, this study does have some limitations. Including a group of BMP-2-treated femora in the bone defect study would have been beneficial in comparing the quality of the regenerated bone between BMP-2 and FK506. The rabbit spine fusion study was limited by the range of doses tested, lacked decorticated and untreated controls, and lacked histological analysis. Additionally, groups were not fully randomized, as the BMP and highest FK506 dose (145mg) were performed bilaterally, while the lower two FK506 doses (64.3mg and 42.6mg) were performed unilaterally. In the future, including a control group that receives a sham decortication procedure but no osteoinductive treatment, adding lower doses of FK506, and performing histological analysis would provide further insight into the efficacy of FK506 as a therapeutic to induce spinal fusion. In addition, since the hMSC work demonstrated the added benefit of combining BMP-2 with FK506, a treatment combining the two would inform the possibility of utilizing FK506 to potentiate a lower dose of BMP-2 required for fusion. This could maximize osteogenic effect without requiring the current supra-physiological BMP-2 dose that currently has several adverse, off-target effects.

These studies build upon previous FK506 research and demonstrates the ability of local FK506 treatment to induce osteogenic differentiation in human cells and bone formation in two pre-clinical *in vivo* orthotopic bone repair models. This is the first demonstration of an osteoinductive small molecule used as a standalone treatment for a critically bone defect or a spinal fusion procedure. In order to translate this therapy towards clinical utilization, future studies will scale local FK506 treatment and test the efficacy in a non-human primate spinal fusion model. This research has the potential to improve healing outcomes for patients and reduce complications caused by current treatments.

## Materials and Methods

### Cell culture

Commercially available primary adult human mesenchymal stem cells (hMSCs) from Donor 47506 were used (Lonza Group Ltd., Basel, Switzerland). Cells were cultured at passage 3 and seeded in triplicate in hMSC Growth Medium (MSCGM™ Mesenchymal Stem Cell Growth Medium, Lonza) with 10% FBS, 1% p/s, and 1% L-Glutamine (Lonza). At 24 hr post seeding, the medium was changed to osteogenic differentiation medium (hMSC Osteogenic Differentiation BulletKit™ Medium, Lonza) with L-ascorbic acid-2-phosphate (AA2P) (5mg/ml stock, 10 µl/ml), and β-glycerolphosphate (BGP) (20 mM stock, 20 µl/ml) (Lonza). The cell culture medium was changed every 3 days. Cultures on various days were subjected to either alkaline phosphatase or mineralization assays.

### Alkaline Phosphatase (ALP) Assay

hMSC were plated at 15,000 cells/well in 24-well plates and grown overnight in a growth medium containing 10% FBS (Lonza). When cells reach 80% confluency, cells were treated with various concentrations of BMP-2 or FK506 in an osteogenic media (Lonza). Cell culture medium was changed every 3 days. On day 12, the cells were washed with phosphate-buffered saline (PBS) and lysed by addition of lysis buffer (10 mM Tris–HCl pH 8.0, 1 mM MgCl2, and 0.5% Triton X-100). The cell lysates were centrifuged for 5 min at 13,000 g. The supernatant was removed and the aliquots were assayed for ALP activity and protein amount. The ALP activity was measured in triplicate using an ALP assay kit (Sigma-Aldrich, St. Louis, MO) in microtiter plates. The protein amount was determined with Bio-Rad protein assay reagent (Bio-Rad, Hercules, CA) using bovine serum albumin (BSA) as a standard. The ALP activity (nmoles of p-nitrophenol per ml) was normalized to the protein amount (nmoles of p-nitrophenol per μg).

### Alizarin Red Staining

hMSC were plated at 15,000 cells/well in 24-well plates and grown overnight in a growth medium containing 10% FBS (Lonza). Once cells reached 80% confluency, they were treated with various concentrations of BMP-2 or FK506 in an osteogenic media (Lonza). The medium was replaced every 3-4 days, and deposition of mineral was observed after 3 weeks. To assess mineralization, the cultures were washed with phosphate-buffered saline and fixed in a solution of ice-cold 70% ethanol for 2-3 h. The cultures were rinsed with water and stained for 10 min with 1 ml of 40 mM alizarin red (pH 4.1). The cultures were rinsed two or three times with phosphate-buffered saline to reduce nonspecific staining, air-dried, and photographed. To assess relative levels of matrix mineralization the Alizarin Red stain was extracted from the samples (3 samples per group) by adding 400 µL of 10% acetic acid followed by 30 min incubation at room temperature. The absorbance at 405 nm of the solubilized Alizarin red dye from the samples was measured using a microplate plate reader (Molecular Devices). An Alizarin red staining standard curve was established with a known concentration of the dye.

### Segmental defect surgery

10-week-old Sprague-Dawley rats underwent a unilateral 5 mm segmental bone defect in the mid diaphysis of the left femur, as previously described ^31^. Briefly, an anterior incision followed by blunt dissection exposed the left femurs. An internal polysulfone fixation plate was fixed to the femurs, and a critically sized 5mm defect was created using Gigli wire saw (RISystem, Davos, Switzerland). Animals were randomly assigned to one of two groups: treatment with FK506 or negative control group. A collagen sponge was used as a vehicle to carry either 4.2 mg of FK506 in 100 µl of Dimethyl sulfoxide (DMSO), or 100 µl of DMSO alone for the negative control group and was implanted into the defect. Bi-weekly radiographs were acquired for a period of 12 weeks at which point the rodents were euthanized and femurs were resected. All femurs underwent µCT analysis and were then used for either histology (n=2/group) or mechanical testing (n=5/group). After surgery, animals were monitored for signs of distress and evaluated for ability to bear weight on the surgical limb during 3-day post-surgery period. One animal from the negative control group had hardware fixation failure and was excluded from the study. All procedures were approved by the Atlanta Veterans Affairs Medical Center IACUC.

### Rabbit spine fusion surgery

6-8 month old female New Zealand White rabbits received single level posterolateral intertransverse process lumbar spine arthrodesis of the 5th and 6th lumbar vertebrae via the bilateral paraspinal muscle-splitting approach as previously described ^32^. Briefly, after the paraspinal muscles were bluntly divided in line with the incision, the transverse processes were exposed and decorticated with an electric burr. Collagen sponge was then implanted based on the treatment groups, and the fascia and skin were then closed with sutures. Bilateral or unilateral procedures were performed and fusion sites were assigned to treatment with one of 4 treatment groups: 1.29 mg BMP-2 in 3 mls PBS (n=4), 145 mg FK506 in 3 mls DMSO (n=4), 64.3 mg FK506 in 2 mls DMSO (n=3), or 42.6 mg FK506 in 2 mls DMSO (n=3). Treatments were loaded onto sterile collagen sponges (two strips/treatment at 15 mm x 40 mm x 3.5 mm for 3 mls DMSO, 15 mm x 30 mm x 3.5 mm for 2 mls DMSO) and implanted. Rabbits were weighed throughout study to monitor weight loss. Rabbits who lost >25% of their baseline body weight were euthanized according to endpoint criteria approved in the IACUC protocol’s Euthanasia Guidelines. All four rabbits that received the bilateral 1.29 mg BMP-2 and 145 mg FK506 treatments exhibited 25% weight loss and were euthanized at 3 weeks, 5.2 weeks, and the last two were euthanized at 6 weeks. One rabbit that received 64.3 mg FK506 exhibited 25% weight loss and was euthanized at week 4. Otherwise, rabbits were euthanized 6 weeks after implantation at which point the spine was resected for µCT analysis. All procedures were approved by the Atlanta Veterans Affairs Medical Center IACUC.

### Micro-computed tomography

Micro-computed tomography (µCT) scans were performed ex vivo (Micro-CT40, Scanco Medical, Bruttisellen, Switzerland). Rat femora were scanned with at 36 µM voxel size at a voltage of 70 kVp and current of 114 µA. Evaluations were performed within defect region on newly formed bone. Rabbit spines were scanned with a 30 µM voxel size at a voltage of 55kVp and current of 145 µA. Scans were contoured to isolate newly formed bone at the fusion site for evaluation.

### Mechanical testing

At 12 weeks, rat hindlimbs were dissected, and fixation plates and soft tissue were removed. Femora were then wrapped in saline-soaked gauze and stored at −80°C until testing. The day of testing, femora were thawed and each end was potted in Wood’s metal (Alfa Aesar). Failure in torsion was tested using a TA Electroforce 3220 load frame at a rotation of 3°/second. Maximum torque to failure was defined as the peak torque value at failure and rigidity was calculated by finding the slope of the linear region of the torque-rotation curve and multiplying that by the gauge length.

### Histology

Rat femurs were fixed in 10% NBF and decalcified with Cal-Ex^®^ II (Fisher Chemical CS5114D Cal-Ex^®^ II Fixative/Decalcifier). Following decalcification and fixation, samples were dehydrated in increasing concentrations of alcohol (70%, 95%, 100%), followed by xylene before embedding in paraffin and cut into 5 micron slices (Accu-Cut SRM 200 Rotary Microtomoe, Sakura Finetek USA, Torrance, CA, USA). Slides were stained with Hematoxylin and Eosin, Goldner’s Trichrome, and Safranin-O/Fast-green. Images were obtained with Aperio VERSA200 (Leica Biosystems, Inc., Buffalo Grove, IL, USA) with a 40X HC PL APO objective with .95 numerical aperture and captured with Aperio ImageScope v12.4.3 (Leica Biosystems, Inc., Buffalo Grove, IL, USA).

### Statistical Analysis

Data are presented as the mean ± standard error of the mean. Results were analyzed using a one-way analysis of variance (ANOVA) with Tukey post-hoc analysis for pairwise comparisons (95% confidence interval) or unpaired t test using GraphPad Prism 9 (GraphPad Prism Software Inc., San Diego, CA, USA).

## Acknowledgements

The authors thank Mila Friedman for performing histological preparation and staining and BioRender for providing the platform to create the graphical abstract. This work was supported in part by: NIH R21 TR001751; NIH R01 AR078892; Veteran Affairs Career Development Award (IK2-BX003845); and the Wu Tsai Human Performance Alliance and the Joe and Clara Tsai Foundation.

## Contributions

JAH: conceptualization, data curation, formal analysis, interpretation, writing, editing

TMF: conceptualization, data curation, formal analysis, editing

SS: conceptualization, data curation, formal analysis, funding acquisition, editing

JK: conceptualization, interpretation, editing

EJD: conceptualization, data curation, editing

CO: data curation, editing

SMP: conceptualization, interpretation, funding acquisition, editing

SDB: conceptualization, interpretation, funding acquisition, editing

NJW: conceptualization, interpretation, funding acquisition, writing, editing

