## Supplemental Table 1 and Supplemental Table 2 for "Local FK506 delivery induces osteogenesis in *in vivo* rat bone defect and rabbit spine fusion models"

**Supplementary Information**

Table S1. Tukey’s Multiple Comparison’s for Alkaline Phosphatase Activity

| Tukey's multiple comparisons test | Mean Diff. | 95.00% CI of diff. | Adjusted P Value |
|---|---|---|---|
| Growth Media vs. 0 ng BMP-2 | -0.1210 | -0.2248 to -0.01721 | 0.0130 |
| Growth Media vs. 40 ng BMP-2 | -0.5233 | -0.6271 to -0.4195 | <0.0001 |
| Growth Media vs. 60 ng BMP-2 | -1.826 | -1.930 to -1.722 | <0.0001 |
| Growth Media vs. 80 ng BMP-2 | -2.582 | -2.686 to -2.478 | <0.0001 |
| Growth Media vs. 160 ng BMP-2 | -5.871 | -5.975 to -5.768 | <0.0001 |
| Growth Media vs. 240 ng BMP-2 | -8.101 | -8.205 to -7.998 | <0.0001 |
| Growth Media vs. 0 µM FK506 | -0.05584 | -0.1596 to 0.04798 | 0.7256 |
| Growth Media vs. 1.25 µM FK506 | -0.4111 | -0.5149 to -0.3073 | <0.0001 |
| Growth Media vs. 2.5 µM FK506 | -0.9367 | -1.041 to -0.8329 | <0.0001 |
| Growth Media vs. 5.0 µM FK506 | -2.098 | -2.202 to -1.994 | <0.0001 |
| Growth Media vs. 10 µM FK506 | -4.265 | -4.369 to -4.161 | <0.0001 |
| 0 ng BMP-2 vs. 40 ng BMP-2 | -0.4023 | -0.5061 to -0.2985 | <0.0001 |
| 0 ng BMP-2 vs. 60 ng BMP-2 | -1.705 | -1.808 to -1.601 | <0.0001 |
| 0 ng BMP-2 vs. 80 ng BMP-2 | -2.461 | -2.565 to -2.357 | <0.0001 |
| 0 ng BMP-2 vs. 160 ng BMP-2 | -5.750 | -5.854 to -5.647 | <0.0001 |
| 0 ng BMP-2 vs. 240 ng BMP-2 | -7.980 | -8.084 to -7.877 | <0.0001 |
| 0 ng BMP-2 vs. 0 µM FK506 | 0.06518 | -0.03863 to 0.1690 | 0.5244 |
| 0 ng BMP-2 vs. 1.25 µM FK506 | -0.2901 | -0.3939 to -0.1863 | <0.0001 |
| 0 ng BMP-2 vs. 2.5 µM FK506 | -0.8157 | -0.9195 to -0.7119 | <0.0001 |
| 0 ng BMP-2 vs. 5.0 µM FK506 | -1.977 | -2.081 to -1.873 | <0.0001 |
| 0 ng BMP-2 vs. 10 µM FK506 | -4.144 | -4.248 to -4.040 | <0.0001 |
| 40 ng BMP-2 vs. 60 ng BMP-2 | -1.302 | -1.406 to -1.199 | <0.0001 |
| 40 ng BMP-2 vs. 80 ng BMP-2 | -2.059 | -2.163 to -1.955 | <0.0001 |
| 40 ng BMP-2 vs. 160 ng BMP-2 | -5.348 | -5.452 to -5.244 | <0.0001 |
| 40 ng BMP-2 vs. 240 ng BMP-2 | -7.578 | -7.682 to -7.474 | <0.0001 |
| 40 ng BMP-2 vs. 0 µM FK506 | 0.4675 | 0.3637 to 0.5713 | <0.0001 |
| 40 ng BMP-2 vs. 1.25 µM FK506 | 0.1122 | 0.008417 to 0.2160 | 0.0262 |
| 40 ng BMP-2 vs. 2.5 µM FK506 | -0.4134 | -0.5172 to -0.3096 | <0.0001 |
| 40 ng BMP-2 vs. 5.0 µM FK506 | -1.575 | -1.679 to -1.471 | <0.0001 |
| 40 ng BMP-2 vs. 10 µM FK506 | -3.742 | -3.845 to -3.638 | <0.0001 |
| 60 ng BMP-2 vs. 80 ng BMP-2 | -0.7565 | -0.8603 to -0.6527 | <0.0001 |
| 60 ng BMP-2 vs. 160 ng BMP-2 | -4.046 | -4.149 to -3.942 | <0.0001 |
| 60 ng BMP-2 vs. 240 ng BMP-2 | -6.276 | -6.380 to -6.172 | <0.0001 |
| 60 ng BMP-2 vs. 0 µM FK506 | 1.770 | 1.666 to 1.874 | <0.0001 |
| 60 ng BMP-2 vs. 1.25 µM FK506 | 1.415 | 1.311 to 1.518 | <0.0001 |
| 60 ng BMP-2 vs. 2.5 µM FK506 | 0.8890 | 0.7852 to 0.9928 | <0.0001 |
| 60 ng BMP-2 vs. 5.0 µM FK506 | -0.2726 | -0.3764 to -0.1688 | <0.0001 |
| 60 ng BMP-2 vs. 10 µM FK506 | -2.439 | -2.543 to -2.335 | <0.0001 |
| 80 ng BMP-2 vs. 160 ng BMP-2 | -3.289 | -3.393 to -3.185 | <0.0001 |
| 80 ng BMP-2 vs. 240 ng BMP-2 | -5.519 | -5.623 to -5.415 | <0.0001 |
| 80 ng BMP-2 vs. 0 µM FK506 | 2.526 | 2.423 to 2.630 | <0.0001 |
| 80 ng BMP-2 vs. 1.25 µM FK506 | 2.171 | 2.067 to 2.275 | <0.0001 |

| | | | |
|---|---|---|---|
| 80 ng BMP-2 vs. 2.5 µM FK506 | 1.645 | 1.542 to 1.749 | <0.0001 |
| 80 ng BMP-2 vs. 5.0 µM FK506 | 0.4839 | 0.3801 to 0.5877 | <0.0001 |
| 80 ng BMP-2 vs. 10 µM FK506 | -1.683 | -1.786 to -1.579 | <0.0001 |
| 160 ng BMP-2 vs. 240 ng BMP-2 | -2.230 | -2.334 to -2.126 | <0.0001 |
| 160 ng BMP-2 vs. 0 µM FK506 | 5.816 | 5.712 to 5.919 | <0.0001 |
| 160 ng BMP-2 vs. 1.25 µM FK506 | 5.460 | 5.356 to 5.564 | <0.0001 |
| 160 ng BMP-2 vs. 2.5 µM FK506 | 4.935 | 4.831 to 5.038 | <0.0001 |
| 160 ng BMP-2 vs. 5.0 µM FK506 | 3.773 | 3.669 to 3.877 | <0.0001 |
| 160 ng BMP-2 vs. 10 µM FK506 | 1.607 | 1.503 to 1.710 | <0.0001 |
| 240 ng BMP-2 vs. 0 µM FK506 | 8.046 | 7.942 to 8.149 | <0.0001 |
| 240 ng BMP-2 vs. 1.25 µM FK506 | 7.690 | 7.586 to 7.794 | <0.0001 |
| 240 ng BMP-2 vs. 2.5 µM FK506 | 7.165 | 7.061 to 7.268 | <0.0001 |
| 240 ng BMP-2 vs. 5.0 µM FK506 | 6.003 | 5.899 to 6.107 | <0.0001 |
| 240 ng BMP-2 vs. 10 µM FK506 | 3.837 | 3.733 to 3.940 | <0.0001 |
| 0 µM FK506 vs. 1.25 µM FK506 | -0.3553 | -0.4591 to -0.2515 | <0.0001 |
| 0 µM FK506 vs. 2.5 µM FK506 | -0.8809 | -0.9847 to -0.7771 | <0.0001 |
| 0 µM FK506 vs. 5.0 µM FK506 | -2.042 | -2.146 to -1.939 | <0.0001 |
| 0 µM FK506 vs. 10 µM FK506 | -4.209 | -4.313 to -4.105 | <0.0001 |
| 1.25 µM FK506 vs. 2.5 µM FK506 | -0.5256 | -0.6294 to -0.4218 | <0.0001 |
| 1.25 µM FK506 vs. 5.0 µM FK506 | -1.687 | -1.791 to -1.583 | <0.0001 |
| 1.25 µM FK506 vs. 10 µM FK506 | -3.854 | -3.958 to -3.750 | <0.0001 |
| 2.5 µM FK506 vs. 5.0 µM FK506 | -1.162 | -1.265 to -1.058 | <0.0001 |
| 2.5 µM FK506 vs. 10 µM FK506 | -3.328 | -3.432 to -3.224 | <0.0001 |
| 5.0 µM FK506 vs. 10 µM FK506 | -2.167 | -2.270 to -2.063 | <0.0001 |

Table S2. Tukey's Multiple Comparison's for Alizarin Red Quantification

| Tukey's multiple comparisons test | Mean Diff. | 95.00% CI of diff. | Adjusted P Value |
|---|---|---|---|
| Growth Media vs. 0 ng BMP-2 | 0.004000 | -0.07760 to 0.08560 | >0.9999 |
| Growth Media vs. 40 ng BMP-2 | -0.02167 | -0.1033 to 0.05993 | 0.9974 |
| Growth Media vs. 60 ng BMP-2 | -0.06333 | -0.1449 to 0.01826 | 0.2412 |
| Growth Media vs. 80 ng BMP-2 | -0.1627 | -0.2443 to -0.08107 | <0.0001 |
| Growth Media vs. 160 ng BMP-2 | -0.3537 | -0.4353 to -0.2721 | <0.0001 |
| Growth Media vs. 240 ng BMP-2 | -0.3400 | -0.4216 to -0.2584 | <0.0001 |
| Growth Media vs. 0 µM FK506 | -0.004667 | -0.08626 to 0.07693 | >0.9999 |
| Growth Media vs. 1.25 µM FK506 | -0.06900 | -0.1506 to 0.01260 | 0.1543 |
| Growth Media vs. 2.5 µM FK506 | -0.1267 | -0.2083 to -0.04507 | 0.0005 |
| Growth Media vs. 5.0 µM FK506 | -0.2117 | -0.2933 to -0.1301 | <0.0001 |
| Growth Media vs. 10 µM FK506 | -0.2553 | -0.3369 to -0.1737 | <0.0001 |
| 0 ng BMP-2 vs. 40 ng BMP-2 | -0.02567 | -0.1073 to 0.05593 | 0.9895 |
| 0 ng BMP-2 vs. 60 ng BMP-2 | -0.06733 | -0.1489 to 0.01426 | 0.1768 |
| 0 ng BMP-2 vs. 80 ng BMP-2 | -0.1667 | -0.2483 to -0.08507 | <0.0001 |
| 0 ng BMP-2 vs. 160 ng BMP-2 | -0.3577 | -0.4393 to -0.2761 | <0.0001 |
| 0 ng BMP-2 vs. 240 ng BMP-2 | -0.3440 | -0.4256 to -0.2624 | <0.0001 |
| 0 ng BMP-2 vs. 0 µM FK506 | -0.008667 | -0.09026 to 0.07293 | >0.9999 |
| 0 ng BMP-2 vs. 1.25 µM FK506 | -0.07300 | -0.1546 to 0.008596 | 0.1098 |

| | | | |
|---|---|---|---|
| 0 ng BMP-2 vs. 2.5 µM FK506 | -0.1307 | -0.2123 to -0.04907 | 0.0003 |
| 0 ng BMP-2 vs. 5.0 µM FK506 | -0.2157 | -0.2973 to -0.1341 | <0.0001 |
| 0 ng BMP-2 vs. 10 µM FK506 | -0.2593 | -0.3409 to -0.1777 | <0.0001 |
| 40 ng BMP-2 vs. 60 ng BMP-2 | -0.04167 | -0.1233 to 0.03993 | 0.7812 |
| 40 ng BMP-2 vs. 80 ng BMP-2 | -0.1410 | -0.2226 to -0.05940 | 0.0001 |
| 40 ng BMP-2 vs. 160 ng BMP-2 | -0.3320 | -0.4136 to -0.2504 | <0.0001 |
| 40 ng BMP-2 vs. 240 ng BMP-2 | -0.3183 | -0.3999 to -0.2367 | <0.0001 |
| 40 ng BMP-2 vs. 0 µM FK506 | 0.01700 | -0.06460 to 0.09860 | 0.9997 |
| 40 ng BMP-2 vs. 1.25 µM FK506 | -0.04733 | -0.1289 to 0.03426 | 0.6327 |
| 40 ng BMP-2 vs. 2.5 µM FK506 | -0.1050 | -0.1866 to -0.02340 | 0.0047 |
| 40 ng BMP-2 vs. 5.0 µM FK506 | -0.1900 | -0.2716 to -0.1084 | <0.0001 |
| 40 ng BMP-2 vs. 10 µM FK506 | -0.2337 | -0.3153 to -0.1521 | <0.0001 |
| 60 ng BMP-2 vs. 80 ng BMP-2 | -0.09933 | -0.1809 to -0.01774 | 0.0084 |
| 60 ng BMP-2 vs. 160 ng BMP-2 | -0.2903 | -0.3719 to -0.2087 | <0.0001 |
| 60 ng BMP-2 vs. 240 ng BMP-2 | -0.2767 | -0.3583 to -0.1951 | <0.0001 |
| 60 ng BMP-2 vs. 0 µM FK506 | 0.05867 | -0.02293 to 0.1403 | 0.3359 |
| 60 ng BMP-2 vs. 1.25 µM FK506 | -0.005667 | -0.08726 to 0.07593 | >0.9999 |
| 60 ng BMP-2 vs. 2.5 µM FK506 | -0.06333 | -0.1449 to 0.01826 | 0.2412 |
| 60 ng BMP-2 vs. 5.0 µM FK506 | -0.1483 | -0.2299 to -0.06674 | <0.0001 |
| 60 ng BMP-2 vs. 10 µM FK506 | -0.1920 | -0.2736 to -0.1104 | <0.0001 |
| 80 ng BMP-2 vs. 160 ng BMP-2 | -0.1910 | -0.2726 to -0.1094 | <0.0001 |
| 80 ng BMP-2 vs. 240 ng BMP-2 | -0.1773 | -0.2589 to -0.09574 | <0.0001 |
| 80 ng BMP-2 vs. 0 µM FK506 | 0.1580 | 0.07640 to 0.2396 | <0.0001 |
| 80 ng BMP-2 vs. 1.25 µM FK506 | 0.09367 | 0.01207 to 0.1753 | 0.0151 |
| 80 ng BMP-2 vs. 2.5 µM FK506 | 0.03600 | -0.04560 to 0.1176 | 0.8963 |
| 80 ng BMP-2 vs. 5.0 µM FK506 | -0.04900 | -0.1306 to 0.03260 | 0.5864 |
| 80 ng BMP-2 vs. 10 µM FK506 | -0.09267 | -0.1743 to -0.01107 | 0.0168 |
| 160 ng BMP-2 vs. 240 ng BMP-2 | 0.01367 | -0.06793 to 0.09526 | >0.9999 |
| 160 ng BMP-2 vs. 0 µM FK506 | 0.3490 | 0.2674 to 0.4306 | <0.0001 |
| 160 ng BMP-2 vs. 1.25 µM FK506 | 0.2847 | 0.2031 to 0.3663 | <0.0001 |
| 160 ng BMP-2 vs. 2.5 µM FK506 | 0.2270 | 0.1454 to 0.3086 | <0.0001 |
| 160 ng BMP-2 vs. 5.0 µM FK506 | 0.1420 | 0.06040 to 0.2236 | <0.0001 |
| 160 ng BMP-2 vs. 10 µM FK506 | 0.09833 | 0.01674 to 0.1799 | 0.0094 |
| 240 ng BMP-2 vs. 0 µM FK506 | 0.3353 | 0.2537 to 0.4169 | <0.0001 |
| 240 ng BMP-2 vs. 1.25 µM FK506 | 0.2710 | 0.1894 to 0.3526 | <0.0001 |
| 240 ng BMP-2 vs. 2.5 µM FK506 | 0.2133 | 0.1317 to 0.2949 | <0.0001 |
| 240 ng BMP-2 vs. 5.0 µM FK506 | 0.1283 | 0.04674 to 0.2099 | 0.0004 |
| 240 ng BMP-2 vs. 10 µM FK506 | 0.08467 | 0.003070 to 0.1663 | 0.0372 |
| 0 µM FK506 vs. 1.25 µM FK506 | -0.06433 | -0.1459 to 0.01726 | 0.2237 |
| 0 µM FK506 vs. 2.5 µM FK506 | -0.1220 | -0.2036 to -0.04040 | 0.0008 |
| 0 µM FK506 vs. 5.0 µM FK506 | -0.2070 | -0.2886 to -0.1254 | <0.0001 |
| 0 µM FK506 vs. 10 µM FK506 | -0.2507 | -0.3323 to -0.1691 | <0.0001 |
| 1.25 µM FK506 vs. 2.5 µM FK506 | -0.05767 | -0.1393 to 0.02393 | 0.3589 |
| 1.25 µM FK506 vs. 5.0 µM FK506 | -0.1427 | -0.2243 to -0.06107 | <0.0001 |
| 1.25 µM FK506 vs. 10 µM FK506 | -0.1863 | -0.2679 to -0.1047 | <0.0001 |
| 2.5 µM FK506 vs. 5.0 µM FK506 | -0.08500 | -0.1666 to -0.003404 | 0.0360 |
| 2.5 µM FK506 vs. 10 µM FK506 | -0.1287 | -0.2103 to -0.04707 | 0.0004 |
| 5.0 µM FK506 vs. 10 µM FK506 | -0.04367 | -0.1253 to 0.03793 | 0.7314 |
